# A content-based representational scaffold for naturalistic event memories

**DOI:** 10.1101/2021.04.16.439894

**Authors:** Zachariah M. Reagh, Charan Ranganath

## Abstract

Although every life event is unique, there are considerable commonalities across events. However, little is known about whether or how the brain flexibly represents information about different event components at encoding and during remembering. Here, we show that different cortico-hippocampal networks systematically represent specific components of events depicted in videos, both during viewing and during episodic memory retrieval. Regions of an Anterior Temporal Network represented information about people, generalizing across contexts, whereas regions of a Posterior Medial Network represented context information, generalizing across people. Medial prefrontal cortex generalized across videos depicting the same schema, whereas the hippocampus maintained event-specific representations. Similar effects were seen in real-time and recall, suggesting reuse of event components across overlapping episodic memories. These findings reveal a computationally optimal strategy in cortico-hippocampal networks for encoding different high-level event components, supporting efficient reuse for event comprehension, imagination, and recollection.

## Introduction

As Heraclitus famously said, you cannot step twice into the same river. The events that make up our lives are extracted from complex, dynamic experiences with pieces that never perfectly repeat. Yet many events have a predictable structure and overlapping event components, such as a particular person, allowing us to generalize across experiences. It can be advantageous for the brain to establish and reuse high-level representations of different event components in order to efficiently form new memories and make inferences about the people, places, and situations that comprise them. However, little is known about whether or how the brain pulls apart and flexibly recombines elements across continuous events, like those in the real world, as they are experienced and recalled.

Recent evidence has indicated that the brain’s ‘default mode network’ (DMN) carries patterns of activity that distinguish one event from another in ongoing narratives, such as television episodes. Based on these studies, neural activity patterns in the DMN seem to reflect a broad understanding of an event, rather than specific elements of that event. For instance, DMN regions carry activity patterns over entire scenes of a movie, which are similar to those evoked when recalling that movie^1–3^. Moreover, activity patterns in DMN regions remain stable as an event unfolds, but these patterns shift abruptly at event transitions^4,5^. Other studies have suggested that certain components of the DMN – particularly, medial prefrontal cortex (mPFC) – might represent abstract information about events that might generalize across instances of similar situations^6^. Related work suggests that DMN regions may carry information related to prior schematic knowledge about television shows and characters in those shows^7^. These findings indicate that the DMN maintains high-level representations that generalize across events. However, the nature of these representations, and the extent to which event content is represented in a unitary or modality-specific way, is not agreed upon.

Much of the DMN is comprised of parietal regions which have been proposed to support retrieval of event content, irrespective of information type or modality^8–11^. These regions have also been proposed to represent generalized semantic information about events^12–14^. Consistent with this view, recent work suggests that DMN regions may uniformly represent all aspects of an event in an integrated manner, with memory strength perhaps teasing apart the contributions of particular regions of the DMN^15^.

An alternative view is that the DMN can be divided into at least two subnetworks that interact with the hippocampus (HPC) to support event cognition and episodic memory^16–18^. According to this view, a Posterior Medial (PM) Network preferentially represents contextual and situational information at multiple levels of abstraction^19^, with parietal and parahippocampal cortex representing information about specific event contexts and mPFC representing schematic information that generalizes across similar situations^20–24^. Conversely, an Anterior Temporal (AT) Network is thought to represent information about entities, such as objects and people^25–27^. Information from the PM and AT Networks, in turn, may be integrated by the hippocampus, enabling information from events to be dissected, flexibly recombined in real time, and reconstructed during episodic memory retrieval.

Although several recent studies have investigated DMN activity during complex, naturalistic events such as films and narratives^28,29^, it is unknown whether the DMN represents individual events in a unitary way, or whether it carries different representations of people, contexts, and situations that can be flexibly constructed and recombined across multiple events. If event representations across the DMN are distributed and content-invariant, we would expect to see uniform representation of event content across the entire DMN, with any nonoverlap across events reducing the similarity of those representations across regions. Conversely, different components of the DMN may act to scaffold different pieces of the experience. Specifically, the PM Network may preferentially represent some components of complex events (contextual information), whereas the AT Network may represent others (entities), and event representations across regions may depend on information content. Prior studies have shown dissociable representations of characters and spatial locations in a movie with divergent narratives^30^ and in imagined autobiographical events^31^. However, importantly, these prior studies did not focus on the stability of representations across multiple events with varying degrees of overlap. That is, we do not know whether some event information can be stably represented and reused in the face of interference from other kinds of information. In other words: are event representations scaffolded in the DMN on the basis of content, and reused across experiences?

Here, we designed a novel set of video clips depicting real-world events, systematically combining information about entities (people) and contexts (four locations). We further manipulated contextual specificity (two classes of contexts with two exemplars of each), allowing us to examine whether any regions of the PM Network – for example, mPFC – would merge across contexts which share abstract similarity. Participants underwent fMRI scanning during encoding and recall of these videos, and multi-voxel activity patterns were analyzed to assess representational similarity across events. In sum, our study merges this level of experimental control with the use of continuous events, allowing us to ask the critical question: are different elements extracted from and stably represented across dynamic situations such as those we encounter in our daily lives?

## Results

Participants viewed 8 35-second video clips combining a central person, each appearing in 1 of 4 contexts (Fig. 1A). To address context specificity or generalization, we incorporated 2 different *types* of contexts (2 distinct cafes and 2 distinct grocery stores). During encoding, participants viewed each event once per run, in randomized order, over 3 total encoding runs (i.e., each video was viewed a total of 3 times). Each clip consisted of a 5-second title screen, followed by the 35-second event, and a 10-second interval between events (Fig. 1B). During the single recall run, participants viewed the title screen for each event for 40 seconds (i.e., no video clip was played), during which they verbally recalled the event in as much detail as possible, followed by a 10-second interval between recall of events (Fig. 1C). Participants showed strong recall and recognition of all events, and showed no evidence of memory biases for or against event content (Table 1). See Methods for additional details, and Supplemental Information for statistical tests pertaining to behavioral performance.

**Figure 1:**
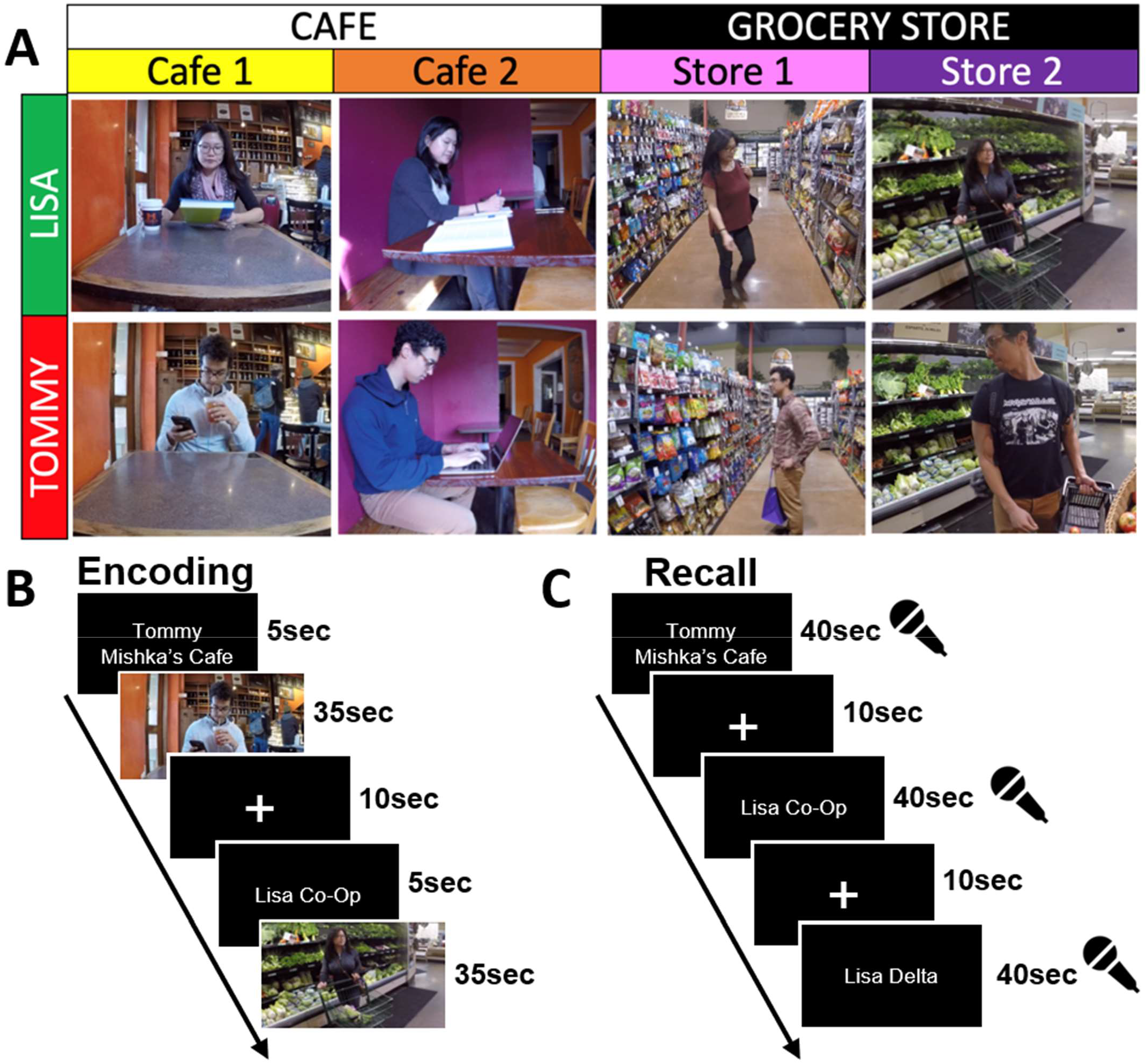
Experimental design and trial structure. (A) 8 videos were designed to systematically combine information about local entities (i.e., central person) and contexts (i.e., specific location), with an additional layer of context *type* (i.e., café vs. grocery store). (B) Encoding trial structure (3 runs, randomized event order). (C) Recall trial structure (1 run, randomized event order).

**Table 1:** Recall and recognition memory performance. Recall (verifiable details, top row) and true/false recognition memory (d’, bottom row) and per event are significantly above zero for every context and person depicted, but do not differ as a function of context or person.

|  | Context |  |  |  | Person |  |
| --- | --- | --- | --- | --- | --- | --- |
|  | Cafe1 | Cafe2 | Store1 | Store2 | Tommy | Lisa |
| **Recall (details)** | 14.893 | 14.225 | 14.317 | 14.601 | 14.773 | 14.594 |
| **Recognition ($d'$)** | 1.493 | 1.445 | 1.457 | 1.518 | 1.733 | 1.704 |

### Regional differences in context, person, schema, and episode-specific patterns at encoding

Briefly, we separately modeled the unique activity pattern for each video clip. To characterize event representations during encoding, we correlated patterns of activity across events and across runs. To test a priori hypotheses about representational content across regions of interest (ROIs; Supplemental Fig. 1), event-by-event correlation matrices for each region were compared to hypothesized model matrices depicting Person, Context, Schema, and Episode-Specific (or Episodic) representations (Fig. 2). Per our experimental design, schemas were operationalized as similarity across a *type* of context (e.g., Cafe1 + Cafe2, or Store1 + Store2). As we expected similar response profiles across PM Network (PMC, ANG, and PHC) and AT Network regions (PRC and TP) (corroborated by analyses over individual ROIs; see Supplemental Information), we collapsed within-network across ROIs. Further details can be found in Methods, and full regional results (including PMC broken into constituent medial parietal subregions, Supplemental Fig. 3) as well as corroborating pattern similarity analyses broken down by event types can be found in Supplemental Information.

**Figure 2:**
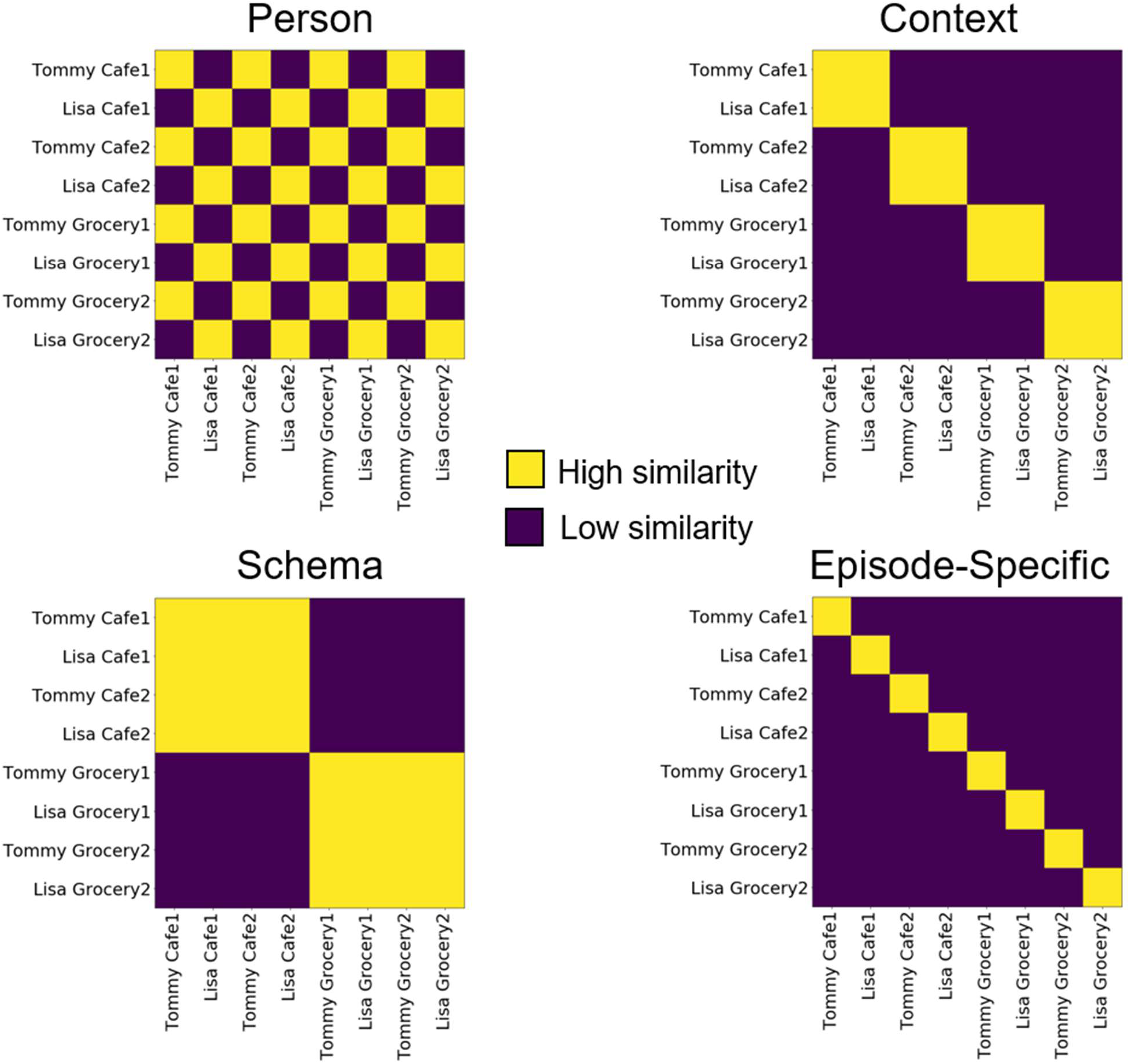
Model matrices based on hypothesized representational profiles. Event-by-event correlation matrices resulting from pattern similarity analyses were compared to hypothesized model matrices depicting Character, Context, Schema, and Episode-Specific representations via point-biserial correlations.

Event-by-event correlation matrices were compared to model matrices to test specific hypothesized representational profiles. In the PM Network (Fig. 3A), the average correlation matrix across participants was best described by the Context model matrix (r = 0.842, p = 2.88e^-18^) with significant correlations also found for the Schema (r = 0.629, p = 2.675e^-08^) and Episodic (r = 0.603, p = 1.326 e^-07^) matrices (Fig. 3B). The Person matrix fit was not significant (r = 0.095, p = 0.453). Model fits differed significantly (F(3,57) = 16.187, p = 9.95e^-08^), and the Context matrix fit was significantly stronger than each of the other three (p_Tukey_ < 0.05 corrected). Follow-up analyses confirmed that similar patterns of results were seen across the different PM Network ROIs (Supplemental Fig. 2).

**Figure 3:**
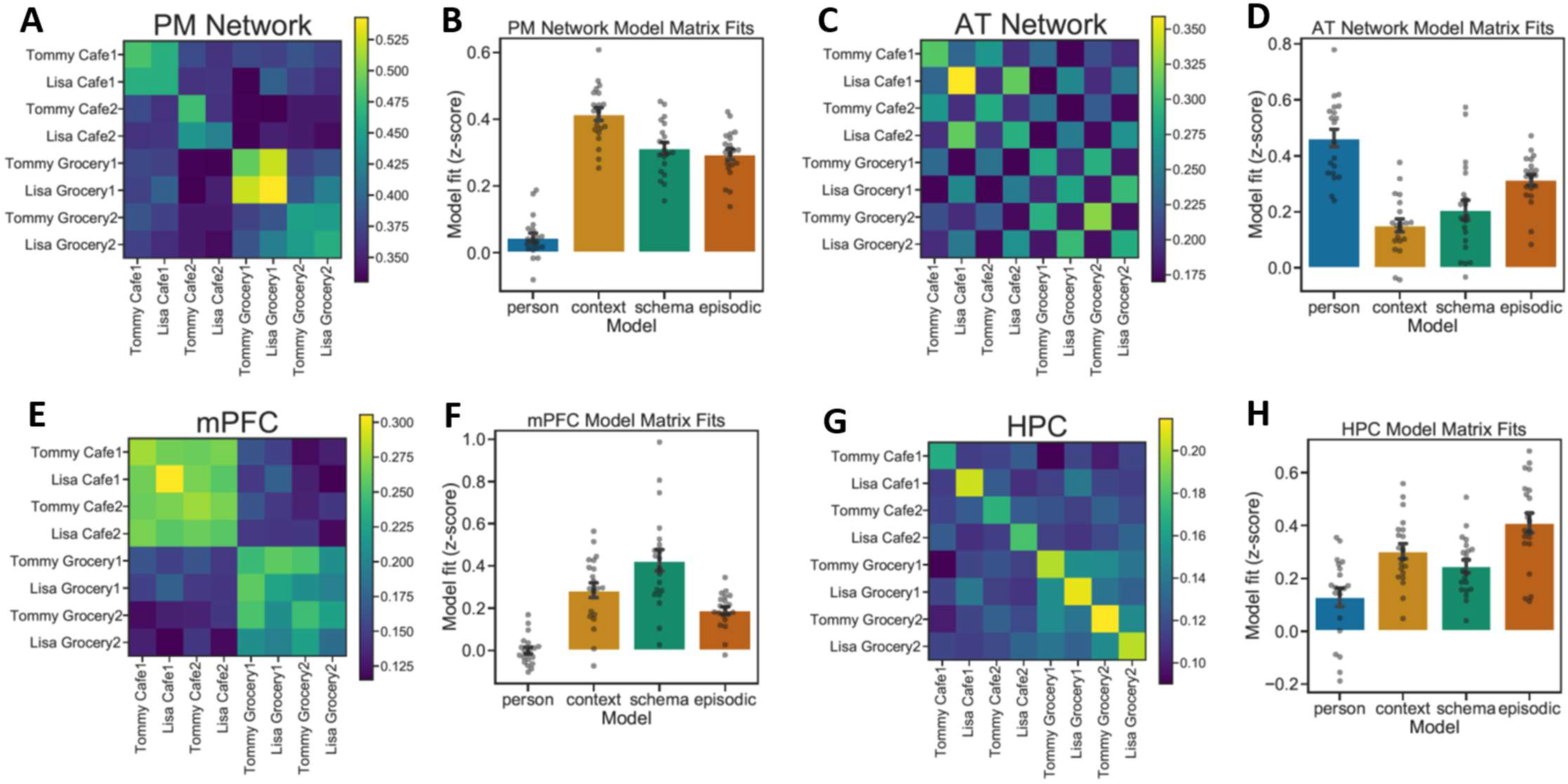
Across-event pattern similarity at encoding. (A,B) PM Network pattern similarity results most strongly fit the Context model matrix. (C,D) AT Network pattern similarity results most strongly fit the Person model matrix. (E,F) mPFC pattern similarity results most strongly fit the Schema model matrix. (G,H) HPC pattern similarity results most strongly fit the Episodic model matrix. (Error bars = SEM.)

In the AT Network (Fig. 3C), the averaged correlation matrix was best described by the Person model matrix (r = 0.845, p = 1.722e^-18^), though fits for all other matrices were also significant: Context (r = 0.274, p = 0.028), Schema (r = 0.366, p = 0.003), Episodic (r = 0.575, p = 6.664e^-07^) (Fig. 3D). Model fits differed significantly (F(3,57) = 18.9, p = 1.24e^-08^), and the Person matrix was a significantly stronger fit than all other model matrices (p_Tukey_ < 0.05 corrected). Like PM Network analyses, AT Network regions (PRC, TP) showed highly similar response profiles (Supplemental Fig. 4).

The averaged correlation matrix in mPFC (Fig. 3E), was best described by the Schema model matrix (r = 0.914, p = 5.089e^-26^), with significant correlations also with the Context (r = 0.0.591, p = 2.759e^-07^) and Episodic (r = 0.385, p = 0.002) matrices (Fig. 3F). The fit with the Person matrix was not significant (r = 0.001, p = 0.995). Model fits differed significantly (F(3,57) = 42.916, p = 1.22e^-14^), with the Schema matrix being a stronger fit than the others (p_Tukey_ < 0.05 corrected).

Event correlations in HPC (Fig. 3G), were best described by the Episodic model matrix (r = 0.588, p = 1.305e^-18^) (Fig. 3H). Significant fits were also observed with the Context (r = 0.67, p = 1.38e^-09^), Schema (r = 0.535, p = 5.377e^-06^), and Person (r = 0.247, p = 0.049) matrices. Model fits differed significantly (F(3,57) = 29.6975, p = 1.08e^-11^). The Episodic model matrix was a stronger fit to the data than the Person or Schema matrices (p_Tukey_ < 0.05 corrected), though there was not a significant pairwise difference between the Episodic and Context matrices.

In contrast to model matrix comparisons, another method of analyzing the pattern similarity results reported in the main text is to collapse across events of a particular kind, and compare to other events of a different kind (e.g., [same character + same context] versus [same character + different context]). Per this analysis, we can factorially cross character (same vs. different) and context (same, similar, or different) and directly contrast z-scored pattern similarity scores across conditions. In the PM Network, we found a significant effect of Context (F(2,114) = 6.325, p = 0.002), with significant effects in each individual region (see Supplemental Information). This effect was driven by greater pattern similarity when participants viewed events that occurred within the same context compared to similar (pairwise contrasts: p_Tukey_ < 0.05 corrected) or different contexts (pairwise contrasts: p_Tukey_ < 0.05 corrected) (Fig. 4A). Pattern similarity did not differ between similar and different contexts. Conversely, we did not find an effect of Person (F(1,114) = 0.24, p = 0.625) nor an interaction (F(2,114) = 0.018, p = 0.982). In the AT Network, we found a significant effect of Person (F(1,114) = 20.139, p = 1.73e^-05^), but no effect of Context (F(2,114) = 1.617, p = 0.203) nor an interaction (F(2,114) = 0.194, p = 0.824). This was driven by significantly greater pattern similarity when participants viewed events depicting the same person compared to a different person (pairwise contrasts: p_Tukey_ < 0.05 corrected) (Fig. 4B). In mPFC, we found a significant effect of Context (F(2,114) = 13.569, p = 5.18e^-06^), but neither an effect of Person (F(1,114) = 0.108, p = 0.742) nor an interaction (F(2,114) = 0.09, p = 0.914). Here, we found higher pattern similarity for same versus different contexts like the PM Network. However, unlike the PM Network, we also found higher pattern similarity for similar versus different contexts (pairwise contrasts: p_Tukey_ < 0.05 corrected), but not between same and similar contexts (Fig. 4C). Finally, in HPC, we found a significant effect of Context (F(2,114) = 5.018, p = 0.008) and a trending interaction (F(2,114) = 2.997, p = 0.054), but no effect of Person (F(1,114) = 1.96, p = 0.164). In line with model matrix analyses, post-hoc contrasts revealed that pattern similarity was highest in HPC when viewing the same event (i.e., same person + same context; pairwise contrasts: p_Tukey_ < 0.05 corrected) (Fig. 4D).

**Figure 4:**
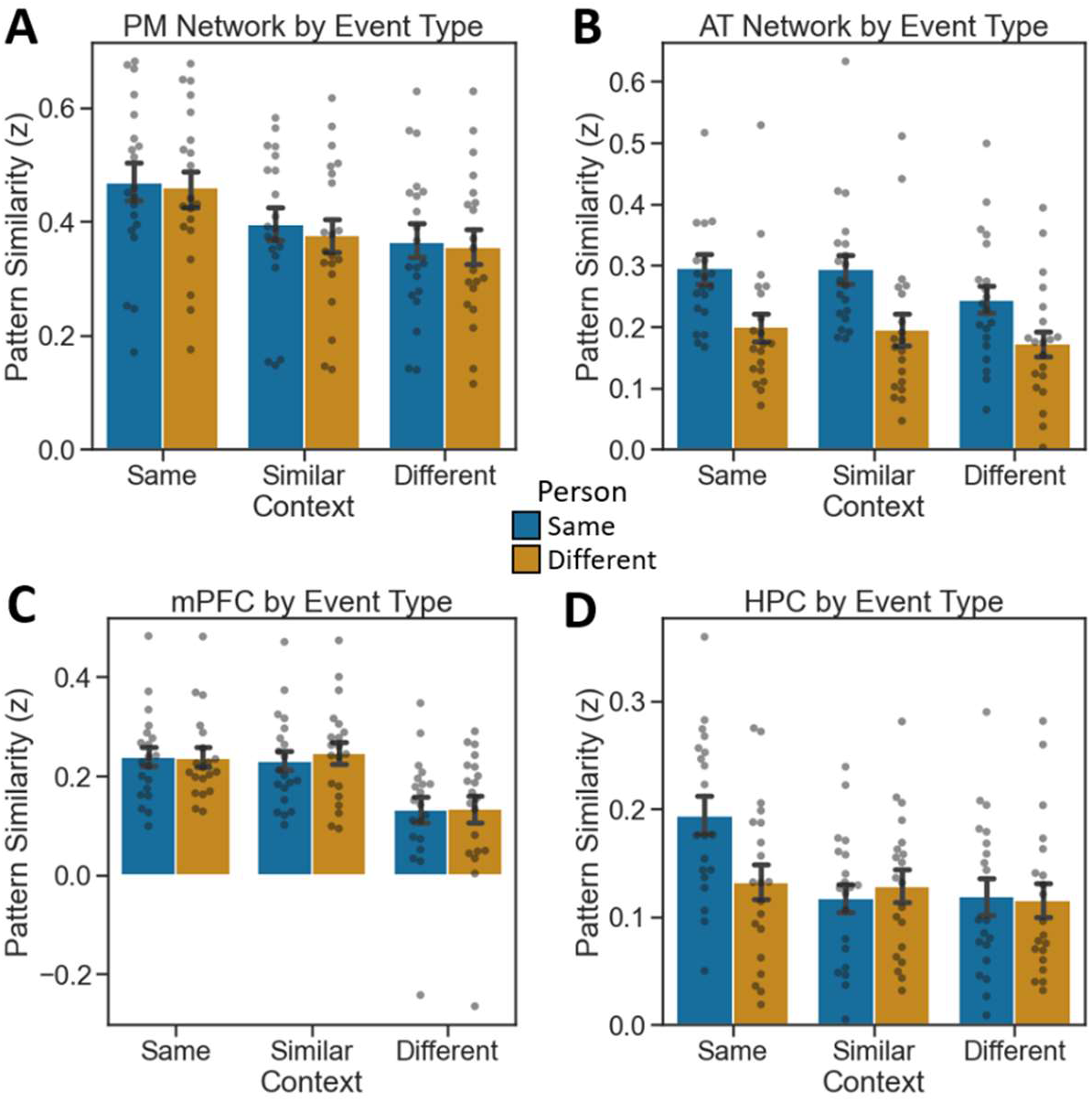
Encoding pattern similarity results by event type. (A) PM Network patterns show context-specificity. (B) AT Network patterns show person-specificity. (C) mPFC patterns generalize across similar contexts. (D) HPC patterns are episode-specific. (Error bars = SEM.)

### Content-selective neural reinstatement of event components during spoken recall

We next asked whether neural patterns associated event content would be reinstated during spoken recall, and whether reinstated patterns would be modality-selective. Whole-event neural patterns were assessed in line with encoding-encoding analyses. For encoding-recall comparisons, we correlated patterns of activity from each event at encoding runs to event patterns for the single recall run, averaged across the three run comparisons. In line with above analyses, we collapsed across PM Network regions and AT Network regions (see Supplemental Information for full regional results), and event-by-event correlational patterns were compared to hypothesized model matrices, resulting in point-biserial correlations assessing model matrix fits (Fig. 2). Further details can be found in Methods.

During recall, the PM Network (Fig. 5A) was best described by the Context model matrix (r = 0.592, p = 2.604e^-07^), though a significant correlation was also observed with the Schema matrix (r = 0.426, p = 0.0004). The fit to the Episodic model was trending, but nonsignificant (r = 0.243, p = 0.053), and there was a significant negative fit to the Person model (r = −0.458, p = 0.0001). Model fits differed significantly (F(3,57) = 20.475, p = 3.97e^-09^), driven by a poorer fit to the Person matrix than other matrices (p_Tukey_ < 0.05 corrected) (Fig. 5B). Though the Context matrix was numerically the strongest fit to PMN reinstatement patterns at recall, this fit did not differ statistically from Schema and Episodic matrix fits at corrected thresholds.

**Figure 5:**
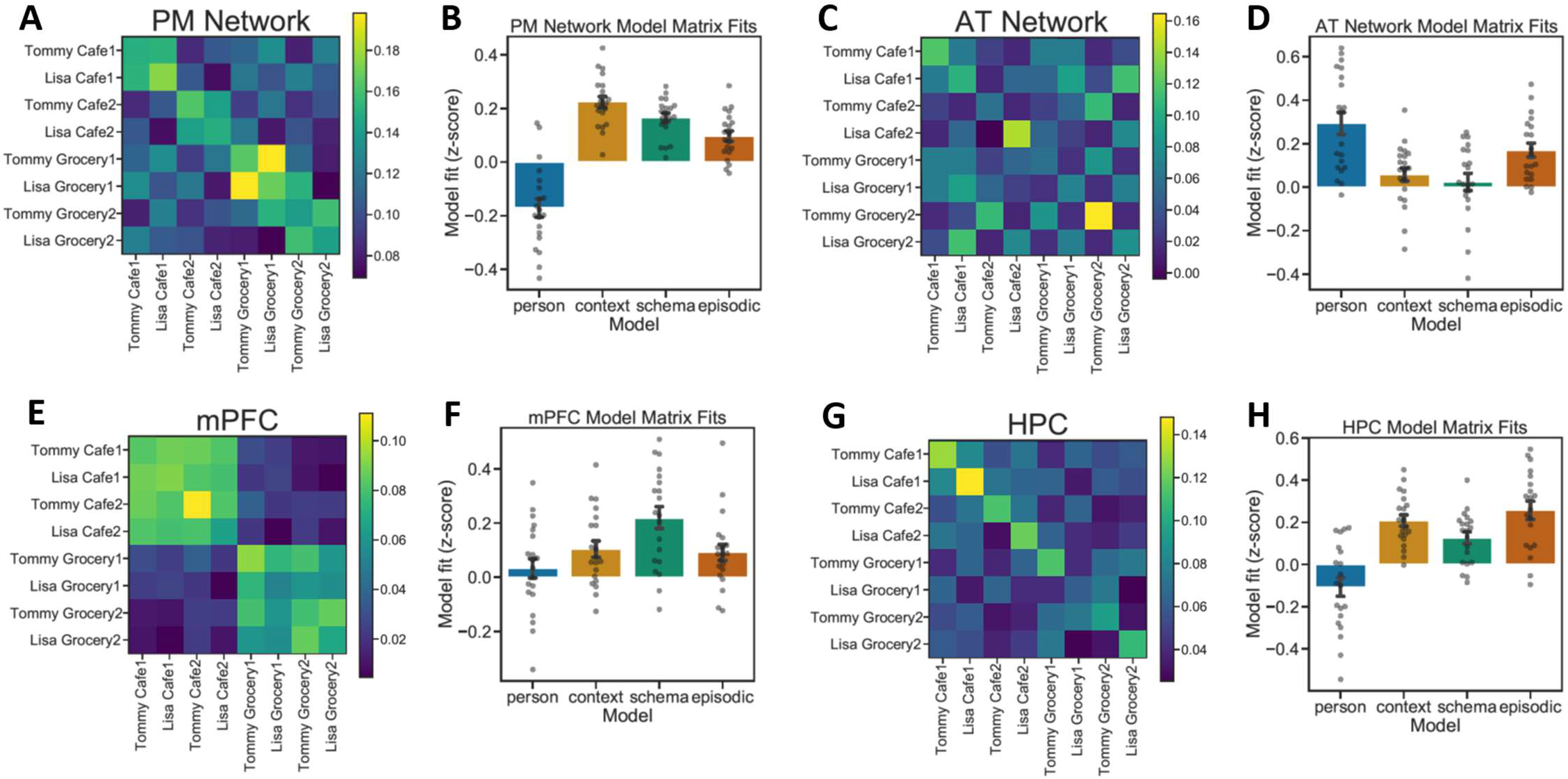
Content-selective pattern reinstatement across ROIs during spoken recall. (A,B) PM Network pattern reinstatement most strongly fits the Context model matrix. (C,D) AT Network pattern reinstatement most strongly fits the Person model matrix. (E,F) mPFC pattern reinstatement most strongly fits the Schema model matrix. (G,H) HPC pattern reinstatement most strongly fits the Episodic model matrix. (Error bars = SEM.)

Encoding-retrieval pattern similarity in the AT Network (Fig. 5C), was best described by the Person matrix (r = 0.539, p = 2.904e^-06^), with a significant fit also to the Episodic matrix (r = 0.307, p = 0.014). The AT Network did not show significant correlations with the Context (r = 0.053, p = 0.677) or Schema (r = 0.144, p = 0.255) matrices. Model fits differed significantly (F(3,57) = 12.625, p = 1.95e^-06^), driven by a significantly better fit between the Person matrix and the Context and Schema matrices (p_Tukey_ < 0.05 corrected) (Fig. 5D).

In mPFC (Fig. 5E), encoding-recall pattern similarity data were best described by the Schema matrix (r = 0.774, p = 5.258e^-14^), with significant fits also for the Context (r = 0.424, p = 0.0001) and Episodic matrices (r = 0.369, p = 0.003). The Person matrix fit was not significant (r = 0.008, p = 0.944). Model fits differed significantly (F(3,57) = 7.276, p = 3.25e^-04^), driven by significantly stronger fit between the data and the Schema matrix than other model matrices (p_Tukey_ < 0.05 corrected) (Fig. 5F).

HPC (Fig. 5G) encoding-retrieval pattern was best described by the Episodic matrix (r = 0.615, p = 6.548e^-08^), though fits to all 3 other model matrices were also significant: Context (r = 0.521, p = 1.035e^-05^); Person (r = 0.299, p = 0.016); Schema (r = 0.321, p = 0.009). There was a significant different among model fits (F(3,57) = 27.727, p = 3.459e^-11^). The Episodic matrix was a significantly stronger fit to the data than the Schema or Person matrices (p_Tukey_ < 0.05 corrected), though the Episodic and Context matrices did not differ significantly from one another (Fig. 5H).

Similar to our approach at encoding above, we analyzed recall-related pattern reinstatement as a function of event type. In the PM Network, we found a significant effect of Context (F(2,114) = 3.887, p = 0.023), with significant effects in PMC and ANG, but not PHC (see Supplemental Fig. 5) (Fig. 6A). There was neither a significant effect of Person (F(1,114) = 0.382, p = 0.536), nor an interaction (F(2,114) = 0.005, p = 0.995). The AT Network showed a significant effect of Person (F(1,114) = 4.844, p = 0.029), which was found individually in PRC, but not TP (see Supplemental Fig. 6). We observed no effect of Context (F(2,114) = 0.016, p = 0.984) nor an interaction (F(2,114) = 0.029, p = 0.971) in the AT Network (Fig. 6B). In mPFC, we found a significant effect of Context (F(2,114) = 5.455, p = 0.005), but neither an effect of Person (F(1,114) = 0.441, p = 0.508) nor an interaction (F(2,114) = 0.007, p = 0.993) (Fig. 6C). Finally, HPC did not show a significant effect of either Person (F(1,114) = 2.273, p = 0.134) or Context (F(2,114) = 1.164, p = 0.316), but did show a significant interaction (F(2,114) = 3.597, p = 0.031) (Fig. 6D). Reinstated patterns were weaker and individual mean differences less pronounced than comparisons across encoding epochs, and no post-hoc pairwise contrasts were significant at corrected thresholds

**Figure 6:**
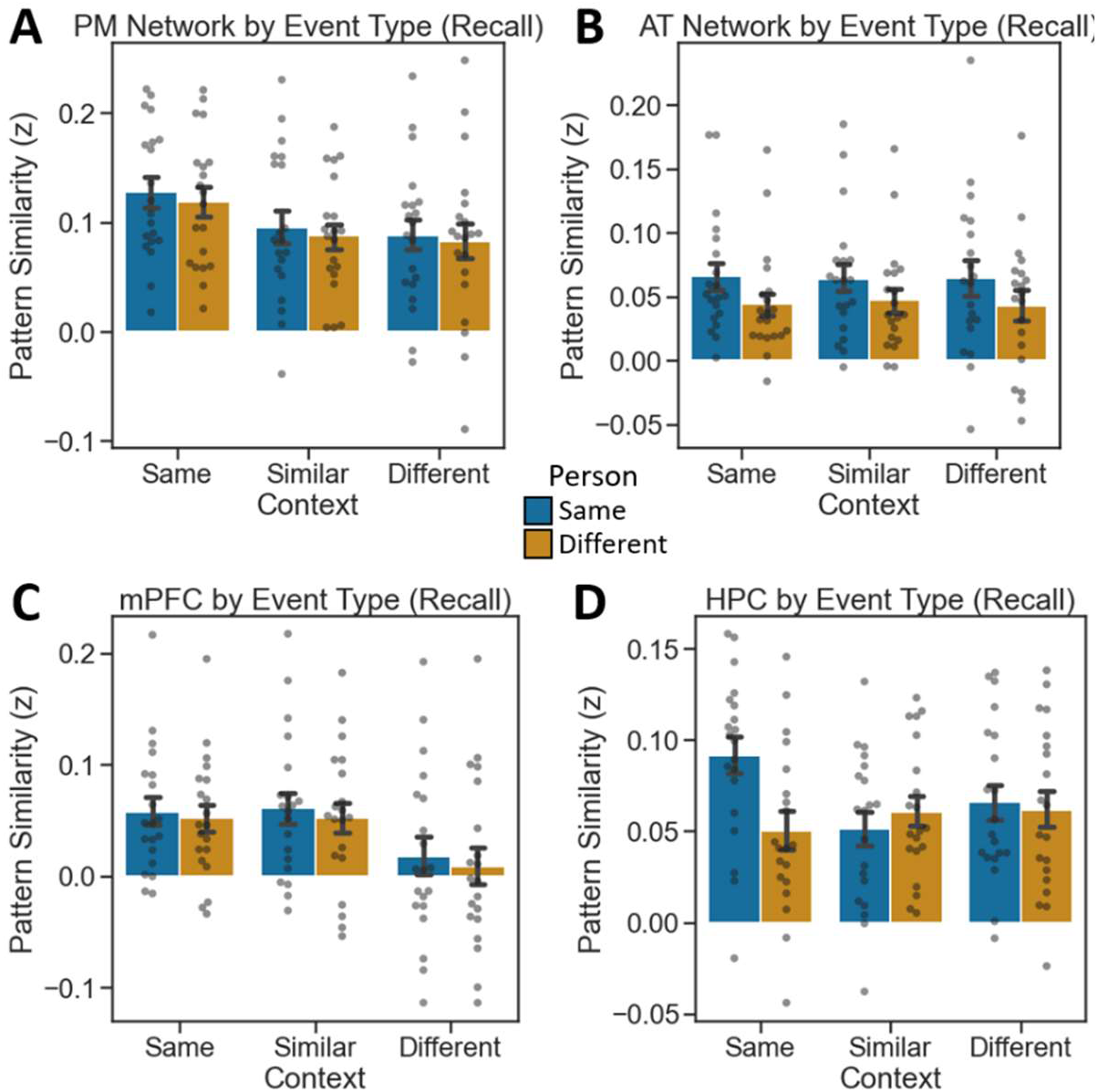
Recall pattern similarity by event type. (A) PM Network patterns show context-specificity. (B) AT Network patterns show person-specificity. (C) mPFC patterns generalize across similar contexts. (D) HPC patterns are episode-specific. (Error bars = SEM.)

### Event pattern reinstatement in the hippocampus correlates with details recalled

Finally, we asked whether neural pattern reinstatement related to recall success. Prior studies have shown that encoding-retrieval similarity, particularly in or mediated by the HPC, correlates with memory performance^32–35^. In the present data, this analysis was conducted by correlating each subject’s HPC encoding-recall pattern similarity, averaged across all same-event comparisons, with the overall number of verifiable details they recalled across all events in the experiment. This was contrasted against the same analysis conducted over all mismatched events rather than same events (i.e., the average pattern similarity value of each event at encoding compared to different events at recall). In other words, this analysis examines the relationship between general HPC pattern reinstatement during recall and overall recall success, comparing matched to mismatched events.

We found that the extent of pattern similarity between encoding and recall for the same events (i.e., encoding-recall match) was significantly correlated with the total number of verifiable details retrieved for all events across participants (r = 0.555, p = 0.011) (Fig. 7A). In contrast, events that were mismatched between encoding and recall showed no evidence of a relationship with HPC reinstatement (r = 0.015, p = 0.475) (Fig. 7B). A Fisher r-to-z transformation reveals a trending (but not statistically significant) difference between the correlations when directly compared via two-tailed test (z = 1.78, p = 0.0751). Though prior studies have found a relationship between reinstatement in PMC and subsequent memory measures (Bird et al., 2015), this relationship was only trending and not significant in the current dataset (r = 0.385, p = 0.094). Moreover, no other ROIs in our data showed a significant relationship with recall. Taken together, these results indicate that neural patterns associated with complex events across multiple encoding episodes are, to some extent, brought back online during recall, and that this reinstatement of neural patterns follows information content dissociations present during encoding. Furthermore, the relationship between encoding and retrieval patterns for specific events in the HPC is correlated with overall recall success. We note that these findings were present when examining the total number of verifiable details across all events. When restricting analyses to reinstatement and recall of specific events, correlations trended similarly but were not statistically significant. Thus, these findings speak to overall relationships between HPC reinstatement and recall success, but cannot speak to this relationship at the level of single events.

**Figure 7:**
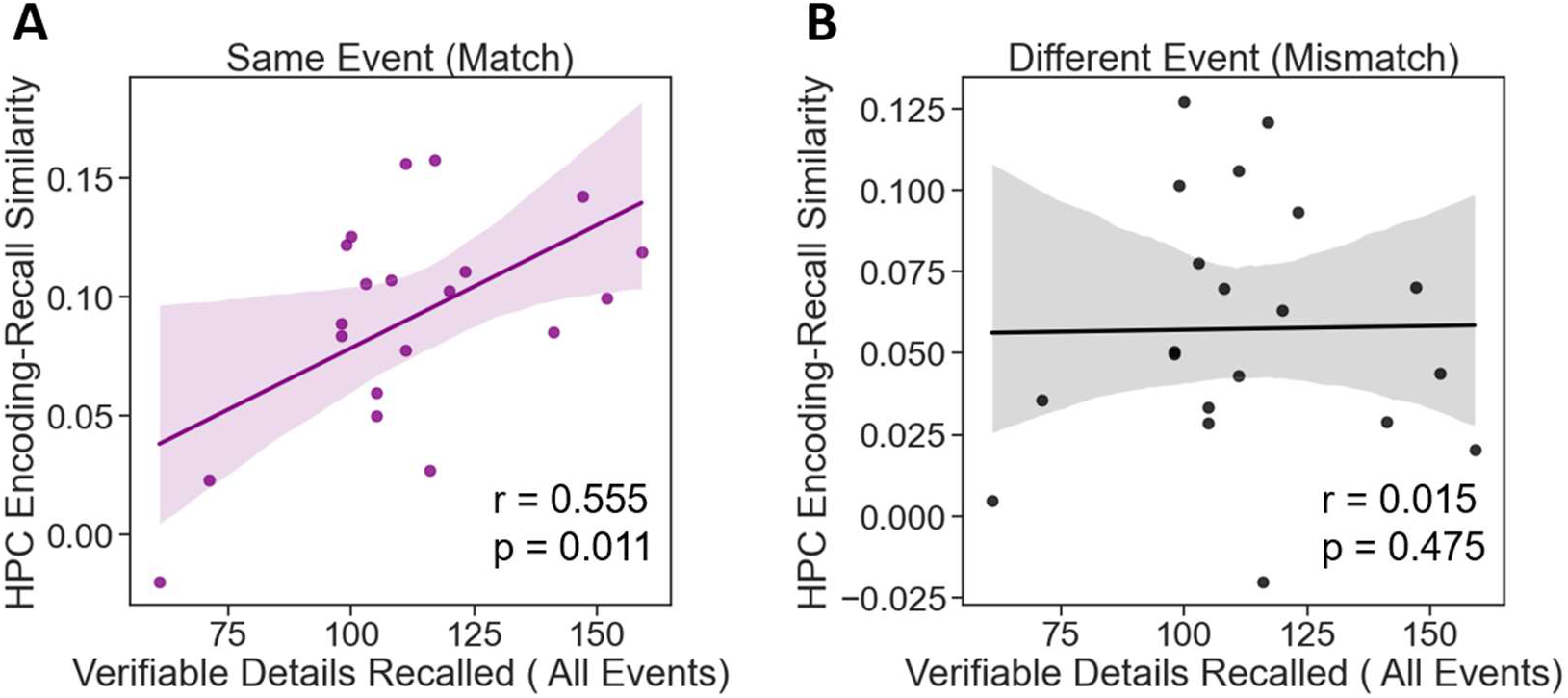
Correlation between HPC reinstatement at recall and the overall number of verifiable details retrieved across all events. Correlations are shown for (A) matched events and (B) mismatched events.

### Distinct timescales of pattern similarity across regions for specific events

In the above analyses, multi-voxel patterns were averaged across the duration of each clip, similar to other recent approaches^1–3^. This was done to capture stable information content being represented across the duration of distinct events. However, we next conducted an exploratory analysis to test whether multi-voxel pattern information during events differed across regions as a function of event epoch. Prior studies suggest distinct temporal receptive windows across the brain, with ‘slow’ timescale regions shifting their activity patterns on the order of tens of seconds to minutes during ongoing experiences^28,36–39^. In these studies, DMN regions have among the slowest representational timescales, which are sensitive to narrative understanding^40^ and have been shown to match human judgments of event transitions^4^.

We compared identical events across runs (i.e., same person, same context) given that all ROIs showed pattern similarity increases during these comparisons but not any other conditions. For these analyses, we modeled each TR in the run individually, in line with beta series analysis^41^. Each TR from a given event in one encoding run was correlated with its corresponding TR in that same event in different runs. We binned these TRs into 3 different eochs, ignoring the first 4 TRs as these featured the event title. The remaining TRs were binned unevenly into event Onset (the first 7 TRs), event Offset (the last 7 TRs), and Mid event (the intervening 15 TRs) epochs. We binned unevenly, opting for relatively shorter Onset and Offset epochs given prior evidence for the importance of HPC activity at event boundaries^42,43^. For each ROI, each epoch was contrasted against its mean pattern similarity value across all events, across all runs (i.e., the grand-mean).

In the PMN (Fig. 8A), pattern similarity for the same event across runs was significantly above the grand-mean at Mid event (t(19) = 2.723, p = 0.014) and event Offset (t(19) = 2.903, p = 0.009), but not at event Onset (t(19) = 1.518, p = 0.145). However, epochs did not differ significantly (F(2,38) = 0.877, p = 0.42). In the ATN (Fig. 8B), pattern similarity was significantly above the grand-mean at event Onset (t(19) = 2.115, p = 0.048) and Mid event (t(19) = 4.916, p = 9.594e^-05^), but not at event Offset (t(19) = 1.269, p = 0.219). Event epochs differed significantly (F(2,38) = 4.903, p = 0.013), and Mid event pattern similarity was greater than event Offset (p_Tukey_ < 0.05 corrected). In mPFC (Fig. 8C), pattern similarity was significantly above the grand-mean at Mid event (t(19) = 3.672, p = 0.002) and event Offset (t(19) = 4.427, p = 0.0002), but not event Onset (t(19) = 1.611, p = 0.124). Event epochs did not differ significantly (F(2,18) = 1.753, p = 0.187). In HPC (Fig. 8D), pattern similarity was significantly different from the grand-mean at event Onset (t(19) = 7.983, p = 1.729e^-07^) and event Offset (t(19) = 5.079, p = 6.656e^-05^), but not at the Mid event epoch (t(19) = 1.491, p = 0.153). Event epochs differed significantly (F(2,38) = 18.836, p = 2.07e^-06^), driven by greater pattern similarity at event Onset than Mid or Offset epochs (p_Tukey_ < 0.05 corrected). Thus, while all ROIs showed increases in pattern similarity for same-event comparisons, there were regional differences in when during the event correlated representations emerged.

**Figure 8:**
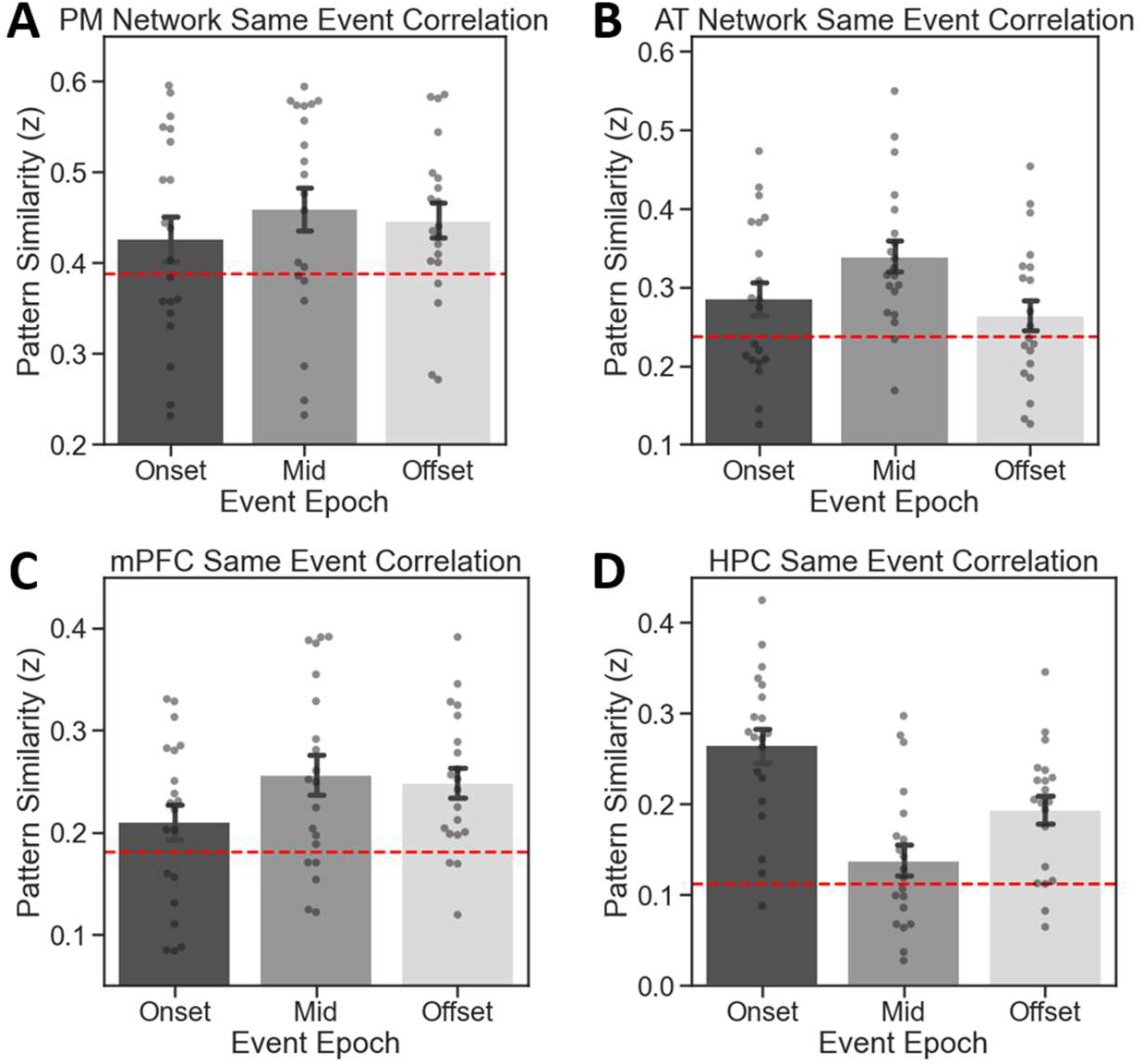
Pattern similarity for same-event comparisons by event epoch. (A) PM Network, (B) AT Network, (C) mPFC, (D) HPC. Red lines indicate the mean pattern similarity value across all events and across all runs for that ROI. (Error bars = SEM.)

## Discussion

Recent research has strongly implicated the DMN as being important for the ongoing perception of and memory for naturalistic events^1–3^. Here, we show that this network systematically deconstructs different aspects of events, such that they can be recombined and reused across different people, places, and situations. We found that areas in the PM Network carried information that generalized across different events that occurred in the same context, and to some extent across events depicting the same general situation. In mPFC, voxel pattern information generalized to a remarkable extent across different events depicting the same situation, consistent with processing of event schemas^6^.

The AT Network, in contrast, carried information about individual people that generalized across films in different contexts and situations. HPC patterns were highly event-specific, in contrast with what was observed in cortical regions, consistent with a role in detailed episodic memory. At recall, HPC pattern reinstatement correlated with retrieved verifiable details about events. Finally, we show evidence for distinct representational timescales for a given event, with HPC being unique in preferentially representing information about event onsets and offsets, but not mid-event epochs.

Previous studies have suggested that the DMN – in particular the posterior medial cortex and angular gyrus – plays an important role in memory, though there are a number of apparently conflicting accounts. One commonly held view is that parietal DMN areas represent retrieved event content in an integrated manner^8–11^. Others have argued that DMN areas represent semantic knowledge, again generalizing across domains and content^12–14^. Other views have emphasized multisensory integration, memory confidence^44^ or precision^15^, or the integration of information across a long timescale^28,39^. The present results are not entirely consistent with any of these views, and yet they might help to reconcile them. Many typical laboratory experiments use information of a single modality (e.g., objects or words, or simple associations between these), which may cause otherwise heterogeneous brain networks to act in a more unitary fashion. Conversely, even if participants recall precise item-level information in these experiments, this may be accompanied by strong reinstatement of contextual or situation associations, which are very difficult to control or even assess across an experimental sample.

Our results – showing preferential representation of contexts and situations by the PM Network and mPFC and preferential representation of people by the AT Network – better align with accounts suggesting that the DMN can be broken into smaller sub-networks. Based on anatomical and functional considerations^16,17^, there is reason to believe that the PM Network might preferentially represent information about the structure of an event, which is used to generate a mental model to guide comprehension and prediction as an event unfolds. In contrast, the AT Network has been proposed to preferentially represent properties of people and things—the “content” that characterizes a specific event. In this framework, the PM and AT Networks can be thought of as “episodic” in that they represent information that is used as a scaffold to understand and remember events^18^, and they can be thought of as “semantic” in that they carry information that generalizes across overlapping events. In general, our data are highly consistent with the PMAT framework^16–18^, and extend this theoretical view to further validate its predictions in the face of complex, continuous events.

Our use of videos depicting real-world events may have permitted the construction and use of event models – mental representations of characters and situations that are strongly tied to the events depicted on the screen, but which also tap into more abstract processes required to understand events that occur in daily life. When recalling a complex event, reinstatement of information from the PM and AT Networks, transferred from other experiences, can serve as a scaffold to guide recovery of precise information about an event. Interestingly, while event-by-event similarity patterns were still consistent with content-selectivity in PM and AT networks during recall, the specificity of this content selectivity (e.g., PM network for context) was reduced compared to encoding-encoding similarity. This could in part be simply due to noisier comparisons when no visual stimulus is available, but an alternative possibility is that these cortico-hippocampal networks are more integrated during memory retrieval compared to encoding. In line with this, a recent study by Cooper and Ritchey^45^ suggest that greater inter-network connectivity may support the multidimensional nature of episodic retrieval.

Our findings are compatible with ideas proposed by Bartlett^46^ and elaborated on by Addis, Wong, and Schacter^47^ suggesting that information in episodic memory can be flexibly recombined in the service of planning, simulation, and imagination. Although studies of episodic simulation have generally focused on the role of the anterior hippocampus, it is also well established that the entire DMN contributes to the generation of imagined situations and events^48^. In our study, hippocampal representations were event-specific, whereas cortical areas carried information that was recombined across events. It is possible that, when imagining or simulating events, people might start by retrieving specific events (driven by hippocampal retrieval), and the PM and AT Networks might enable retrieved information about people, places, and situations to be recombined in novel ways to construct an elaborate event^49^.

Though the hippocampus is universally agreed to be critical for memory, its exact role is also a matter of ongoing study. One widely held view of the hippocampus is that it is important for creating and maintaining unique representations of specific events, linked to a theoretical role in pattern separation^50–52^. Our results are broadly consistent with this view. Our findings also align with studies showing a relationship between reinstatement of specific hippocampal patterns and retrieval success^33–35^. In contrast to representational specificity, other studies strongly suggest that the hippocampus has a role in genearalization^53–56^. Indeed, the hippocampus has the capacity to support specific pattern-separated representations as well as to support generalization via pattern completion^57^. Though our data more strongly support generalization in neocortical regions than the hippocampus, it is possible that the relative lack of specific instructions during event encoding resulted in a stronger bias to create and use specific hippocampal representations rather than to generalize across similar instances. That is, if individuals were incentivized to generalize across similar events (e.g., strategically collapse across events featuring the character Lisa), hippocampal representations may be driven away from specificity and more toward generalization. This possibility can be tested by future experiments.

Although our data support theories of content-based dissociations of the PM and AT Networks, we do not take this to suggest that all regions of each respective network are doing the same thing. For instance, context specificity may differ across regions affiliated with the PM Network. In line with this, PM Network regions showed a significant fit with the Schema model matrix. This is consistent with other findings from Baldassano and colleagues^6^, who used typical progressions of events through airports and restaurants in movie and audio clips to test shared neural responses across different stories featuring schema-congruent event sequences. Baldassano and colleagues reported schema-level representations not only in mPFC, but across other PM Network regions such as PMC and PHC. Their study differs markedly from our approach in that they sought to examine shared neural patterns across highly distinct events sharing a common schematic theme, whereas our approach was to systematically recombine characters and contexts to interrogate representations of specific event content. However, these findings are highly complementary. Additionally, recent work by Barnett et al. suggests that the DMN may feature up to 4 subnetworks rather than just 2^58^. This study uses an atlas generated by Glasser et al., which features a fairly large number of ROIs^59^. We chose to use a more typical approach to ROI delineation, in line with many prior studies^2–6^. Nonetheless, we did run a simple supplemental analysis based on the Barnett et al., networks, which can be found in the Supplemental Information (Supplemental Fig. 7). The general pattern of results is largely in line with the findings as presented here. Future studies can leverage these network-based analytical developments to parse the DMN at increasing levels of granularity.

Closely related to the topic of event memory is that of event structure. Recent studies using naturalistic stimuli such as films have primarily focused on event boundaries rather than the content of events per se. A finding with increasing support is that the hippocampus and certain PM Network regions are transiently more active at event boundaries^4,5,60^, and may moreover strengthen functional interactions at event boundaries^61^. These phenomena are generally not observed in AT Network regions^60^. The present study was not designed to examine event segmentation, but rather to examine the way specific components may be represented across events. However, some insight may be gained from our results. Coarse event boundaries (which are fairly linked to story-related changes rather than individual actions) are reliably elicited by changes in spatiotemporal context^62–64^. The present data would predict PM Network representations to be key at such moments.

Following from this, exploratory analyses also suggest that event representations may come online at distinct epochs as events unfold. In general, cortical regions tended to carry information about event content most strongly in the middle of those events. Conversely, HPC carried event-specific information at event onset and offset. This suggests that representational timescales may be another important factor which differs across regions, broadly consistent with prior work indicating multiple temporal receptive windows across the cortex in response to stimuli such as movies^28,36–39^. Our results also align with an increasingly-supported idea of HPC being particularly important for encoding information at event boundaries^42,43^.

We note that the event depictions in this study were 40 seconds in duration, limiting analyses of the way different representations of event components may evolve over long timescales. Additionally, we did not systematically vary features of characters such as actions, intentions, and other similar factors which could influence event understanding^65–67^. However, having established what may be core components of flexible event representations, future studies can more closely examine these factors with respect to the content of event representations. Another relevant topic is that of schema congruency, which has been found to affect recognition and recall of information^68–71^. However, few studies manipulating and testing effects of schema congruency per se have done so testing memory for lifelike situations^72^. Further, we suggest that our findings take a step beyond general ideas of schema and gist-based memory by systematically varying key event components, such as people, places, and situations. This allows for concrete, constructive formulations of what might make a memory representation generalized versus specific. An approach such as the one used in the present design could be modified to probe questions about schema and gist, critically manipulating congruency between people, their actions, their contextual associations, and overall situations.

In sum, our results address a key gap in existing studies of event cognition and memory by revealing how the complementary functions of different cortico-hippocampal networks enable to brain to flexibly construct and re-use mental representations of event components across different episodic memories. These coding schemes together amount to an optimal and computationally efficient strategy for reducing complex events into key components, representing commonalities across events while also maintaining specific event representations. This provides a representational scaffold for processing, remembering, and even simulating events as we navigate through real-world experiences.

## Methods

### Participants

Participants were recruited from the University of California, Davis and the Davis, California community via flyer and email advertisement. Participants were paid $25 per hour, and gave written informed consent in accordance with the University of California, Davis Institutional Review Board under an approved protocol. Exclusion criteria included magnetic bodily implants, claustrophobia, a history of major psychiatric or neurological disorders, a concussion in the past 6 months, untreated diabetes or hypertension, current drug or alcohol abuse, left-handedness, age outside a range of 18-35 years, and 4 or fewer hours of sleep on the day of data collection.

We initially collected 24 participants, and excluded 1 due to falling asleep during MRI scanning, 2 due to excessive motion in the MRI scanner, and 1 due to equipment malfunction during data collection. The final sample of 20 participants had a mean age of 23.4 years (SD = 3.1 years), consisting of 13 self-identified females and 7 self-identified males.

### Stimuli

8 video clips were developed for use in the present experiment. These video clips were recorded at locations in Davis, California using a GoPro HERO+LCD camera, at 1080p 60fps resolution. Two members of the UC Davis Dynamic Memory Lab served as central characters. Several ‘takes’ were recorded for each video. Videos were then edited in the GoPro editing software to be 35 seconds in duration, and to include a 5 second title screen with white text against a black background for each event depicted (making each video 40 seconds in total).

Prior to video viewing, participants viewed a series of still images depicting the characters and locations subsequently viewed in the event videos. These data were not analyzed here. Videos were presented in the MRI scanner environment using PsychoPy version 3.1.5, and stimulus onset was synchronized with MRI pulse sequences via fiber optic trigger pulses sent to the stimulus computer. Videos were downsampled to from 1080p to 720p in the experiment to reduce the time taken to load and buffer the files in PsychoPy. After video viewing, participants were cued by on-screen text via PsychoPy to recall the events using an MRI-compatible microphone (see Procedure).

After in-scanner tasks, participants were given a True/False recognition test outside the scanner consisting of a series of single-sentence statements about video content. False statements were designed to probe for accuracy of memory, but were not designed to test fine-grained mnemonic discrimination (i.e., they were not designed to be highly similar to factual events). Single sentences were presented in white text against a black background on a 2015 MacBook Pro 13” laptop.

### Procedure

Participants arrived at the UC Davis Center for Neuroscience, and were escorted into the Dynamic Memory Lab testing space where they were briefed on the experiment, and gave written informed consent. Participants then completed an MRI screening form and demographics questionnaires. Upon successful screening for the experiment, participants were escorted to the Imaging Research Center, and were prepped to undergo scanning.

Scanning took place in a Siemens 3T Skyra with a 32-channel head coil. Participants were fitted with MRI-compatible earbuds with replaceable foam inserts (MRIaudio), and were provided with additional foam padding inside the head coil for hearing protection and to mitigate head motion. Participants were additionally given bodily padding, blankets, and corrective lenses as needed. An MRI-compatible microphone (Optoacoustics FOMRI-III) with bidirectional noise-cancelling was affixed to the head coil, and the receiver (covered by a disposable sanitary pop screen) was positioned over the participant’s mouth. Participants were given a description of strategies to remain still while speaking during functional image acquisition. During MRI data acquisition, an eye tracker was operational to monitor participants’ wakefulness and head motion during spoken recall, but no eye tracking data were recorded.

After testing the earbuds and microphone, high-resolution T1-weighted structural images were acquired using a magnetization prepared rapid acquisition gradient echo (MPRAGE) pulse sequence (FOV = 256mm, image matrix = 256 x 256, sagittal slices = 208, thickness = 1mm). Participants then completed a functional imaging run in which they were shown still images of the central people and contexts subsequently shown in the videos (these data are not analyzed or discussed here). Participants then completed 3 runs in which all 8 event videos were shown, in random order across runs and across participants. Video stimuli (title screen + event) were 40 seconds, with a 10 second inter-stimulus interval displaying a fixation cross. Instructions were to remain still and closely attend to the videos, as memory for the videos would be tested later in the experiment. After the 3 encoding runs were completed, participants completed spoken recall of each event. Event titles were displayed in random order, in white text against a black background for 40 seconds, with a 10 second interstimulus interval displaying a fixation cross. Participants were instructed to begin recalling the named event, in as much detail as possible and in order to the extent possible, and to stop recall either when finished or when the event title transitioned to the fixation cross. We reasoned participants would rarely exceed 40 seconds to recall the events based on pilot behavioral data, though we observed 3 instances across all subjects where recall for a given event was cut off due to a time-out. Functional images were acquired using a multi-band gradient echo planar imaging (EPI) sequence (TR = 1220ms, TE = 24ms, FOV = 192mm, image matrix = 64 x 64, flip angle = 67, multi-band factor = 2, axial slices = 38, voxel size = 3mm isotropic, P >> A phase encoding, AC-PC alignment). A single 4 TR functional scan of reverse phase encoding polarity (A >> P) was acquired for unwarping (see fMRI data preprocessing below).

### Behavioral analyses

In-scanner recall was scored using an adapted version of the Autobiographical Memory Interview^73,74^ (see Cohn-Sheehy et al. for a highly similar adapted approach^75^). Three scorers transcribed the audio recording for each event into text, and segmented the document into meaningful detail units (Z.M.R. and two research assistants). Detail units refers to the smallest possible meaningful unit, and labels were assigned to those details on the basis of their content. These details were then classified as “verifiable” if they referred to a factually accurate piece of information pertaining to the events depicted in the videos, and were not preceded by statements of uncertainty (e.g., “maybe”). Redundant information was not counted. Once scoring was completed, interrater reliability was assessed, and was fairly high across the three raters overall (Pearson r = 0.86), and in terms of details scored as being verifiable (Pearson r = 0.84). Verifiable details were compared across characters and across contexts using one-way ANOVAs, and were correlated with pattern similarity and recall-driven reinstatement using Pearson correlations.

While we were interested in the extent to which participants might show intrusions and swaps of details across events that shared a focal person or context, we found that this was very rare in our sample. Specifically, only 2 participants swapped any details across such events, and in both cases these were misattributions of single action details rather than multiple details, or confusion of two person-context pairings in their entirety. Thus, in the present data, there are too few instances of such intrusions or misattributions to be sufficiently powered to characterize such instances. Relatedly, recall was scored purely in terms of verifiably-correct details, meaning they were compared to specific actions or moments depicted in a given episode. Thus, based on behavioral evidence as scored, recall was assessed in terms of successful retrieval of a particular event. Follow-up analyses investigated the prevalence of what might be regarded as “schematic” details which broadly describe a situation but do not refer to specific actions or moments in a clip (e.g., “the barista runs the credit card”), but such instances were similarly too sparse across the data to conduct thorough analyses. In sum, overall recall was clip-specific in the vast majority of instances.

Out-of-scanner recognition test results for each participant considered distinction of true statements from false statements about the viewed events. To quantify this, we calculated a d’ score for each participant: z(Hit rate) – z(False Alarm rate). Similar to in-scanner recall performance above, recognition memory scores were compared across characters and across contexts using one-way ANOVAs, and standard Pearson correlations were used to assess relationships with neural data across events. The ultimate goal of these analyses was to ensure that recognition and recall of event information were significantly nonzero, and unbiased across the events.

### fMRI data preprocessing

Results included in this manuscript come from preprocessing performed using FMRIPREP version stable^76,77^ (RRID:SCR_016216), a Nipype^78,79^ (RRID:SCR_002502) based tool. Each T1w (T1-weighted) volume was corrected for INU (intensity non-uniformity) using N4BiasFieldCorrection v2.1.0^80^ and skull-stripped using antsBrainExtraction.sh v2.1.0 (using the OASIS template). Brain surfaces were reconstructed using recon-all from FreeSurfer v6.0.1^81^ (RRID:SCR_001847), and the brain mask estimated previously was refined with a custom variation of the method to reconcile ANTs-derived and FreeSurfer-derived segmentations of the cortical gray-matter of Mindboggle^82^ (RRID:SCR_002438). Spatial normalization to the ICBM 152 Nonlinear Asymmetrical template version 2009c^83^ (RRID:SCR_008796) was performed through nonlinear registration with the antsRegistration tool of ANTs v2.1.0^84^ (RRID:SCR_004757), using brain-extracted versions of both T1w volume and template. Brain tissue segmentation of cerebrospinal fluid (CSF), white-matter (WM) and gray-matter (GM) was performed on the brain-extracted T1w using fast^85^ (FSL v5.0.9, RRID:SCR_002823).

Functional data was slice time corrected using 3dTshift from AFNI v16.2.07^86^ (RRID:SCR_005927) and motion corrected using mcflirt (FSL v5.0.9^87^). Distortion correction was performed using an implementation of the TOPUP technique^88^ using 3dQwarp (AFNI v16.2.07^86^). This was followed by co-registration to the corresponding T1w using boundary-based registration^89^ with six degrees of freedom, using bbregister (FreeSurfer v6.0.1). Motion correcting transformations, field distortion correcting warp, BOLD-to-T1w transformation and T1w-to-template (MNI) warp were concatenated and applied in a single step using antsApplyTransforms (ANTs v2.1.0) using Lanczos interpolation.

Physiological noise regressors were extracted applying CompCor^90^. Principal components were estimated for the two CompCor variants: temporal (tCompCor) and anatomical (aCompCor). A mask to exclude signal with cortical origin was obtained by eroding the brain mask, ensuring it only contained subcortical structures. Six tCompCor components were then calculated including only the top 5% variable voxels within that subcortical mask. For aCompCor, six components were calculated within the intersection of the subcortical mask and the union of CSF and WM masks calculated in T1w space, after their projection to the native space of each functional run. Frame-wise displacement^91^ was calculated for each functional run using the implementation of Nipype.

Many internal operations of FMRIPREP use Nilearn^92^ (RRID:SCR_001362), principally within the BOLD-processing workflow. For more details of the pipeline see https://fmriprep.readthedocs.io/en/stable/workflows.html.

### fMRI data analysis – representational similarity analysis

All representational similarity analyses (RSA) were run on unsmoothed native-space functional images after the preprocessing steps described above. Each of the 3 encoding runs and the recall run of the raw data were entered into a general linear model (GLM) in AFNI using the 3dDeconvolve function. Nuisance regressors for linear scanner drift (first order polynomial), head motion (6 directions plus their 6 derivatives), and the first two principal components of a combined white matter and CSF mask (via aCompCor above). Given the relatively long 40 second duration of the ‘trials’ being modeled and the 10 second gap between the videos, a least-squares all (LSA) approach was used, such that each run was modeled to produce beta estimate images for either encoding or recall of each of the 8 events. Though a number of prior studies examining naturalistic encoding and recall have not used GLMs to model data, and have instead used a time-shifted averaging approach, we chose to model the present data as this allowed us to more directly mitigate influences of signal drift, motion, and global nuisance signal in evaluating representational patterns. As events, including title screens, were 40 seconds in duration and the TR for our EPI sequence was 1.22 seconds, there were approximately 33 TRs covering each event. Encoding-encoding analyses simply computed event-by-event pattern similarity across runs, such that each event pattern similarity comparison was the average of three across-run comparisons (i.e., run1 to run2, run1 to run3, run2 to run3). For same-event comparisons, this resulted in 3 correlations across runs, which were averaged. For between-event comparisons, this resulted in 6 correlations across runs, which were averaged (twice the number of same-event comparisons). This is comparable to the split representational dissimilarity matrix approach taken by Henriksson and colleagues^93^. Correlation coefficients were then z-scored prior to model matrix comparisons or event-type analyses. Encoding-recall comparisons were followed the same principle as encoding-encoding comparisons, except the 33-TR events modeled at encoding were compared to variable TR events during recall. Specifically, the length of the recall epoch was defined as the duration of spoken recall for each participant (e.g., if a participant spoke for 18 TRs for a given event, the regressor for that event covered those 18 TRs). Recall betas were then compared to encoding betas across all 3 encoding runs.

Resultant beta images were masked using a region-of-interest (ROI) approach, based on theoretical interest in areas of the Posterior-Medial and Anterior-Temporal Networks. ROIs (Supplemental Fig. 1) of the PM Network – angular gyrus (ANG), posterior-medial cortex (PMC), and parahippocamapal cortex (PHC) – were selected based on theoretical involvement in context representation, and involvement in recent studies of naturalistic events. Given accumulating evidence for schema representation in mPFC, we considered this region separately from the PM Network. ROIs in the AT Network – perirhinal cortex (PRC) and temporal poles (TP) – were selected due to demonstrated sensitivity to item-level information. Hippocampal, parahippocampal, and perirhinal ROIs were derived from in-house hand-traced template images in MNI space^94^, which were warped to each subject’s native space using ANTs. All other ROIs are derived from the FreeSurfer Desikan atlas. Multi-voxel patterns were extracted from each ROI mask and written to CSV files for subsequent analyses using the 3dmaskdump function in AFNI. Voxels with null values in any scanning run were excluded.

For model matrix comparisons (see below), each event pattern was correlated with each other event pattern. For conditional comparisons across RSA results, we collapsed events across 4 key conditions: (1) same event (i.e., person-in-context), (2) same person, different context, (3) same person, similar context, (4) same context, different person), and (5) different person, different context. Pairwise comparisons across each event were conducted within subjects, and resulting correlation coefficients were z-scored. For each ROI, events satisfying each condition were averaged, and conditional averages were statistically compared against one another using two-way ANOVAs, with person and context information as factors. Significant effects were evaluated using Tukey’s HSD for post-hoc comparisons.

### fMRI data analysis – model matrix comparisons

To probe specific hypotheses about representation of person, context, schema, and episodic specificity in our ROIs, we compared event-by-event correlation matrices for pattern similarity data to model matrices. Beta images resulting from the preprocessing steps described above underwent pairwise comparisons across events for each participant and for each ROI, resulting in an event-by-event correlation matrix. This was compared to four model matrices: (1) an episode-specific matrix, in which events were hypothesized to correlate only with themselves, (2) a person matrix, in which events were hypothesized to correlate with those events which featured the same person, (3) a context matrix, in which events were hypothesized to correlate with those events which featured the same context, and (4) a schema matrix, in which events were hypothesized to correlate with those events which featured a *similar* context (e.g., two similar but not identical cafes). Model matrices were constructed simply to feature a value of 1 in hypothesized high-correlation cells, and a value of 0 in hypothesized low-correlation cells. In general, model fit was evaluated using a point-biserial correlation between the pattern similarity matrix and the model matrices for each ROI. Model fits were evaluated in two ways. First, for each ROI, we computed the biserial correlation between group average event-wise pattern similarity data and each of the 4 model matrices using the lower-half of the matrix (i.e., excluding redundant comparisons), yielding overall model matrix correlations per region. Second, to incorporate sensitivity to variability across participants and to directly compare different model fits per region, we assessed the model fits for each participant per ROI. For this approach, each participant’s model fit correlations were z-scored, and correlations were aggregated across participants. Finally, one-way ANOVAs and post-hoc Tukey’s HSD comparisons were used to relate the overall fits of the empirical data to the models, and to directly compare fits across models within each ROI. We note that, with this approach, some model matrices will share mutual information (e.g., the Episodic matrix is a component of all other matrices), but we can nonetheless assess the relative fit of each matrix for a given region.

### fMRI data analysis – time-binned analyses

Additional exploratory analyses were conducted examining pattern similarity across events on the order of TRs. Preprocessed data were modeled distinctly from prior analyses: rather than a beta image being generated for each *event*, a beta image was generated for each *time step* comprising an event (similar to a beta-series correlation approach^41^. Confound regressors were unchanged from the other GLM which modeled whole events with single regressors. The analysis consisted of a TR-by-TR pattern similarity analysis within an event pair (rather than event-by-event pattern similarity analysis) across ROIs and collapsed across participants. Given that different ROIs responded to different conditions, we chose to compare only same-event neural patterns across runs for this analysis, as this is the only condition in which all ROIs reliably showed pattern similarity increases across participants. Furthermore, given evidence for distinct mechanisms of hippocampal encoding at event boundaries^42,43^, we binned the time series to attempt to isolate boundary-relevant activity from mid-event patterns. We ignored the first 4 TRs as these timepoints featured the event title. We binned the next TRs as follows: event Onset (the first 7 TRs), event Offset (the final 7 TRs), and Mid event timepoints (the intermediate 15 TRs). This additionally simplified analyses of the data while nonetheless allowing for evaluation of a temporally-evolving change in pattern similarity. We conducted two sets of analyses for these data. The first analysis was a series of one-sample t-tests against the mean pattern similarity value across all encoding runs, evaluating whether pattern similarity for same-event comparisons was significantly different from the baseline pattern similarity of that ROI across all events, at each of the 3 epochs we defined. The second analysis was a one-way ANOVA with post hoc Tukey’s HSD to test whether the 3 event epochs differed significantly from one another.

## Acknowledgments

We thank Jessica Macaluso and Ryan Bugsch for assisting with scoring of recall transcripts. We thank current and former members of the Dynamic Memory Lab for crucial feedback, with special thanks to Brendan Cohn-Sheehy, Jordan Crivelli-Decker, and Derek Huffman. We thank Alex Barnett and Kamin Kim for acting in the videos in addition to providing helpful input. We thank Sarah Morse for helpful input on the manuscript. We thank Delta of Venus, Mishka’s Café, The Nugget, and Davis Food Co-Op in Davis, California for allowing us to record videos featuring their businesses. Finally, we thank our funding sources supporting this work: ONR Grant N00014-15-1-0033 to C.R. and NIA T32AG050061 to Z.M.R.

## Author Contributions

Z.M.R. and C.R. designed the research and wrote the paper. Z.M.R. conducted the experiments and analyzed the data.

