## Supplemental Information for "A content-based representational scaffold for naturalistic event memories"

### Supplemental Information: A cortico-hippocampal scaffold for representing and recalling lifelike events

#### Supplemental Results

##### Model matrix results for individual ROIs: encoding

In ANG (Supplemental Fig. 2A), the average pattern similarity correlation matrix across participants was most strongly related to the Context model matrix (r = 0.794, p = 5.193e$^{-15}$) with significant correlations also found for the Schema (r = 0.604, p = 1.247e$^{-07}$) and Episodic (r = 0.529, p = 6.881 e$^{-06}$) matrices (Supplemental Fig. 2B). The Person matrix correlation was not significant. There was a significant difference among model fits (F(3,57) = 83.206, p < 0.001), and the Context matrix was a significantly stronger fit to the data than each of the other three ($p_{Tukey}$ < 0.05 corrected). Similarly, in PMC (Supplemental Fig. 2C), pattern similarity data most strongly correlated with the Context matrix (r = 0.856, p = 0.586e$^{-19}$) and significant relationships were also observed compared to the Schema (r = 0.657, p = 3.679e$^{-09}$) and Episodic (r = 0.607, p = 1.048e$^{-07}$) matrices (Supplemental Fig. 2D). The Person matrix correlation was not significant. Model fits significantly differed from one another (F(3,57) = 71.763, p < 0.001), with a stronger Context matrix fit than all others ($p_{Tukey}$ < 0.05 corrected). PHC (Supplemental Fig. 2E) also showed this profile, with significant fits to the Context (r = 0.626, p = 3.235e$^{-08}$), Schema (r = 0.413, p = 0.001), and Episodic (r = 0.521, p = 1.019e$^{-05}$) model matrices, and a poor fit to the Person matrix (r = 0.141, p = 0.266) (Supplemental Fig. 2F). There was a significant difference among model fits (F(3,57) = 16.187, p < 0.001), but only the Context vs. Person matrix fits differed significantly ($p_{Tukey}$ < 0.05 corrected).

In line with recent work, we included a posterior-medial cortex (PMC) ROI comprising medial parietal subregions. This raises questions of whether constituent cortical regions – namely, retrosplenial cortex (RSC), posterior cingulate cortex (PCC), and precuneus show sufficiently similar correlation profiles to warrant collapsing across these regions. We examined event-by-event correlation matrices for these three constituent ROIs, which were very similar individually to the correlation matrix produced by averaging them together into a PMC ROI (Supplemental Fig. 3). Indeed, contrasting correlation coefficients between each of the three correlation matrices and model matrices against the PMC fits, no correlation approached a significant difference (all p > 0.63). This provides support for collapsing these ROIs into PMC, and further supports the notion of medial parietal regions comprising a representationally-coherent PM Network.

In PRC (Supplemental Fig. 4A), we found the strongest correlation between the event-by-event pattern similarity matrix and the Person model matrix (r = 0.838, p = 6.117e$^{-18}$), though all other model matrix fits were significant as well: Context (r = 0.249, p = 0.047), Schema (r = 0.322, p = 0.009), and Episodic (r = 0.551, p = 2.424e$^{-06}$) (Supplemental Fig. 4B). Model fits differed significantly (F(3,57) = 33.42, p < 0.001), and the Person matrix was a significantly stronger fit than all other model matrices ($p_{Tukey}$ < 0.05 corrected). Similar results were observed in TP (Supplemental Fig. 4C), where the Person matrix fit the observed data the strongest (r = 0.0.928, p = 2.628e$^{-17}$), but the other model matrix fits were significant as well: Context (r = 0.295, p = 0.018), Schema (r = 0.41, p = 0.001), and Episodic (r = 0.588, p = 3.301e$^{-07}$) (Supplemental Fig. 4D). There was a significant difference among model fits (F(3,57) = 18.9, p < 0.001), and the Person matrix fit the data significantly better than the Person or Schema matrices ($p_{Tukey}$ < 0.05 corrected), but did not differ in a direct pairwise contrast with the Episodic matrix.

**Model matrix results for individual ROIs: recall**

In ANG (Supplemental Fig. 5A), the average pattern similarity correlation matrix across participants was most strongly related to the Context model matrix (r = 0.619, p = 4.769e$^{-08}$) with significant correlations also found for the Schema (r = 0.401, p = 0.001), Episodic (r = 0.287, p = 0.021) and matrices, and a significant negative fit to the Person matrix (r = -0.274, p = 0.028) (Supplemental Fig. 5B). There was a significant difference among model fits (F(3,57) = 20.4752, p < 0.001), with the Context matrix being a stronger fit than Person or Episodic ($p_{Tukey}$ < 0.05 corrected). In PMC (Supplemental Fig. 5C), we found the strongest correlation between encoding-recall pattern similarity data and the Context matrix (r = 0.635, p = 1.771e$^{-08}$), though significant correlations were also observed with the Schema (r = 0.415, p = 0.001), and Episodic (r = 0.288, p = 021) matrices (Supplemental Fig. 5D). The fit with the Person matrix was significant, but negative (r = -0.359, p = 0.003). Model fits differed significantly (F(3,57) = 38.612, p < 0.001), driven by a stronger fit between the Context matrix and other model matrices ($p_{Tukey}$ < 0.05 corrected). PHC (Supplemental Fig. 5E) showed the largest correlation with the Context matrix (r = 0.402, p = 0.001), and was also significantly correlated with the Schema matrix (r = 0.358, p = 0.004) (Supplemental Fig. 5F). Similar to the prior PM Network regions, PHC showed a significant negative fit to the Person matrix at recall (r = 0.567, p = 1.034e$^{-06}$). Model fits differed significantly (F(3,57) = 55.481, p < 0.001), driven by a significantly poorer fit for the Person matrix than others ($p_{Tukey}$ < 0.05 corrected).

In PRC (Supplemental Fig. 6A), we observed significant correlations only with the Person (r = 0.323, p = 0.009) and Episodic (r = 0.267, p = 0.033) model matrices (Supplemental Fig. 6B). The Context and Schema matrix fits were not significant. Model fits differed significantly from one another (F(3,57) = 2.799, p = 0.048), but pairwise differences were not significant. In TP (Supplemental Fig. 6C), we observed significant correlations with the Person matrix (r = 0.291, p = 0.003) and the Episodic matrix (r = 0.272, p = 0.005) (Supplemental Fig. 6D). The Context and Schema matrix fits were not significant, and model fits did not differ significantly from one another (F(3,57) = 1.689, p = 0.179).

**Supplemental Figures**

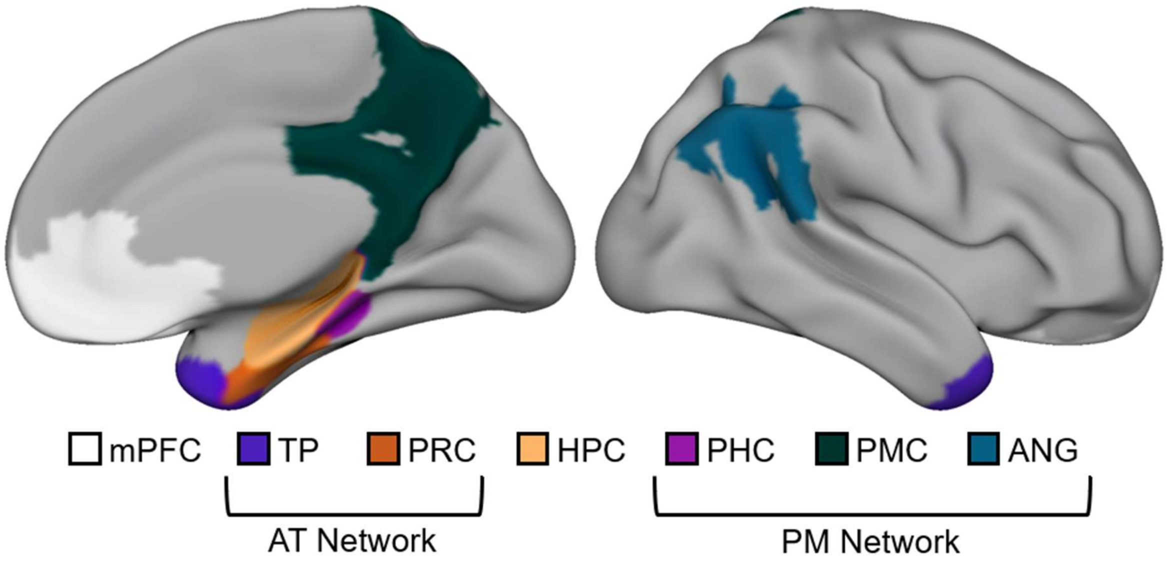


**Supplemental Figure 1: Regions of Interest (ROIs).** Medial temporal lobe ROIs (HPC, PRC, PHC) were adapted from prior anatomical tracings (Ritchey et al., 2015). Cortical ROIs (mPFC, TP, PMC, ANG) were selected from FreeSurfer segmentations. ROIs are displayed on an inflated brain in MNI space using the Surf Ice software package (https://www.nitrc.org/projects/surfice/).

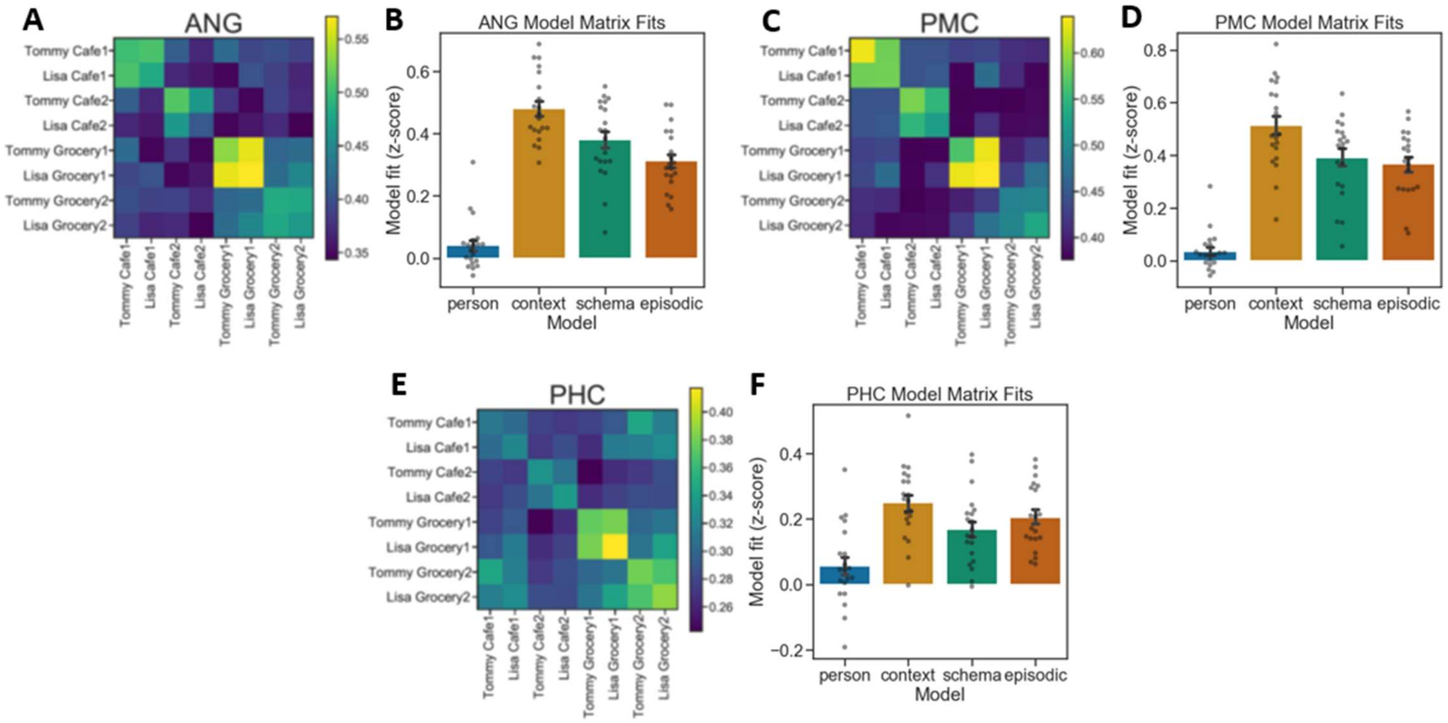


**Supplemental Figure 2: Across-event pattern similarity at encoding for individual PM Network ROIs.** The strongest fit was observed between the Context matrix and across-event pattern similarity data in (A,B) ANG, (C,D) PMC, and (E,F) PHC.

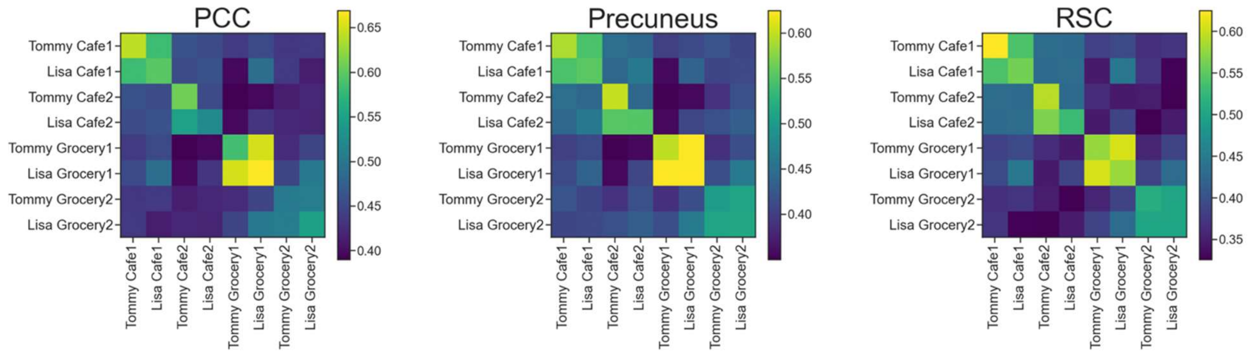


**Supplemental Figure 3: ROIs comprising PMC.** Event-by-event correlation matrices for individual medial parietal ROIs (posterior cingulate cortex: PCC; precuneus, retrosplenial cortex: RSC) comprising our posterior medial cortex (PMC) ROI, as shown below in Supplemental Figure 3. Correlation matrices are qualitatively very similar to PMC, and correlations between each of these 3 ROIs and our model matrices do not differ statistically from fits observed in PMC, as seen above in Supplementa. Fig. 2C,D (all $p > 0.63$).

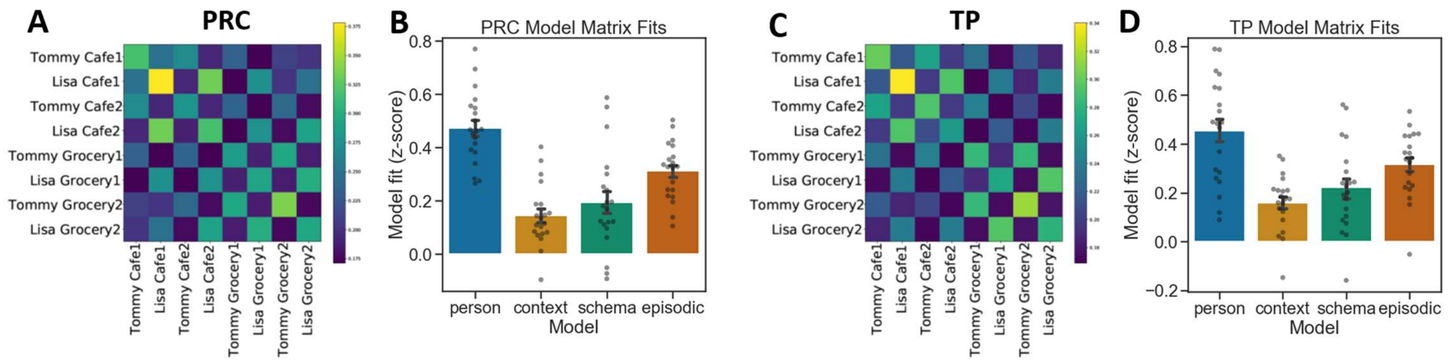


**Supplemental Figure 4: Across-event pattern similarity at encoding for individual AT Network ROIs.** The strongest fit was observed between the Person matrix and across-event pattern similarity data in (A,B) PRC and (C,D) TP.

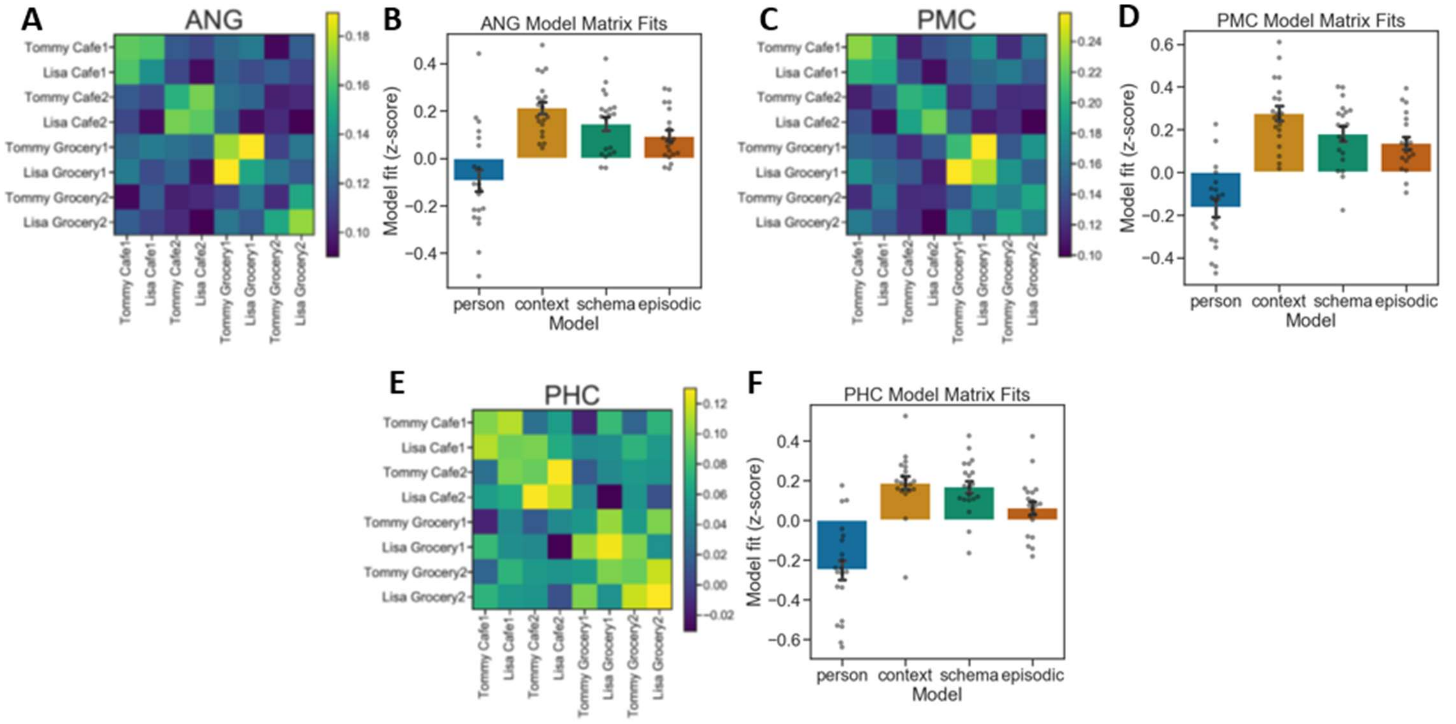


**Supplemental Figure 5: Encoding-recall pattern similarity for individual PM Network ROIs.** The strongest fit was observed between the Context matrix and across-event pattern similarity data in (A,B) ANG, (C,D) PMC, and (E,F) PHC.

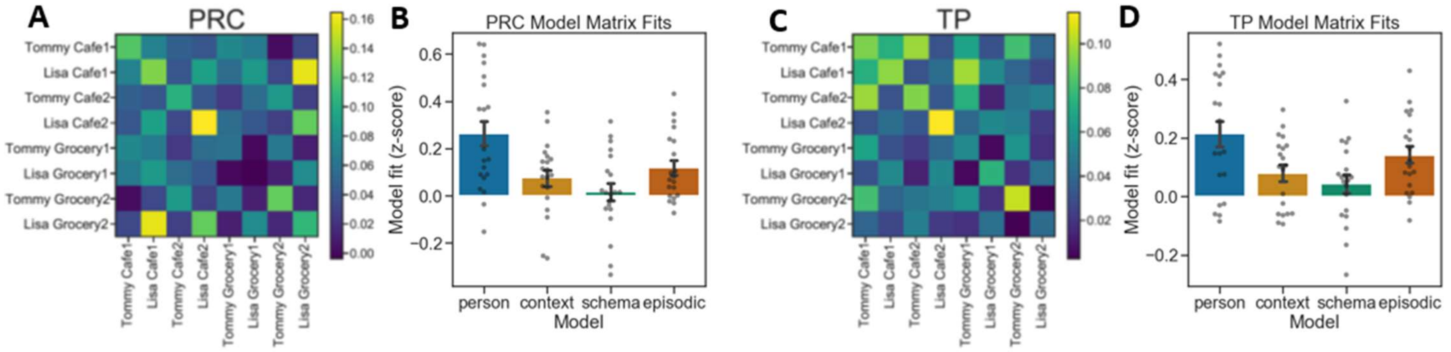


**Supplemental Figure 6: Encoding-recall pattern similarity for individual AT Network ROIs.** The strongest fit was observed between the Person matrix and across-event pattern similarity data in (A,B) PRC and (C,D) TP.

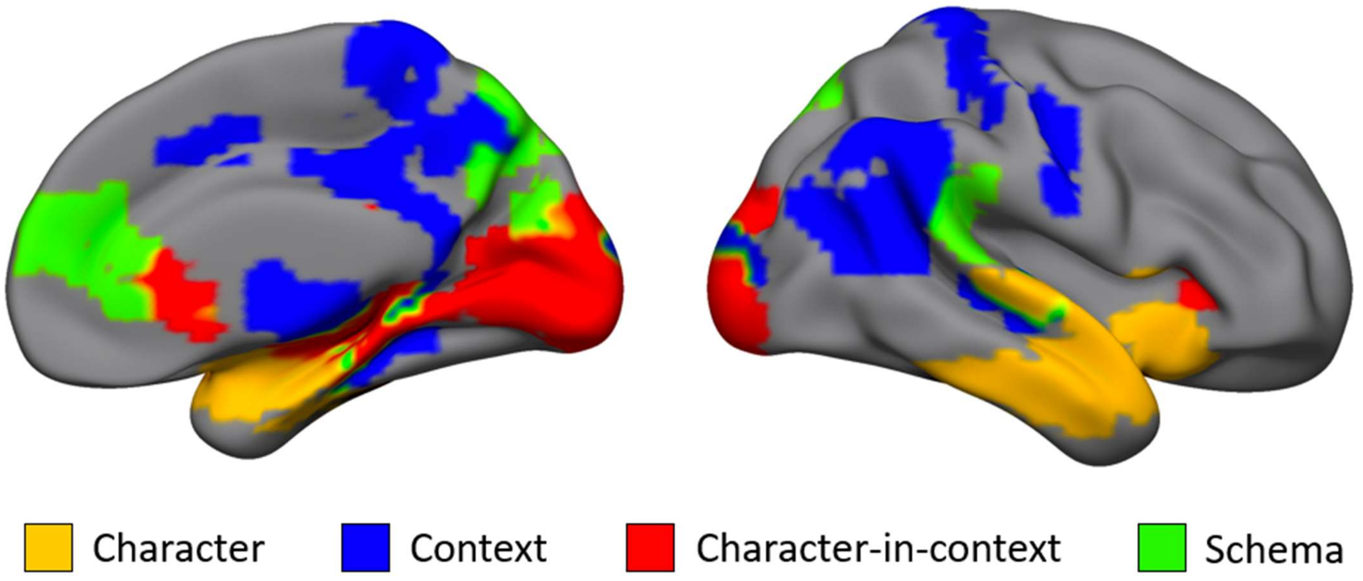


**Supplemental Figure 7: Model matrix fits mapped onto cortical surface regions-of-interest.** An exploratory analysis was conducted at uncorrected thresholds based on Barnett et al., *PLOS Biology* 2021, which identified 4 cortico-hippocampal networks (PM Network, AT Network, a Medial Prefrontal Network, and a Medial Temporal Network). Model matrix comparisons were conducted in each ROI featured in the Barnett et al., atlas (built on the ROIs featured in Glasser et al., *Nature* 2016). A simple "winner-take-all" analyses was conducted in that, if significant model matrix fits were observed, the strongest-fitting effect was plotted in that ROI. Findings are generally consistent with the more limited ROI-based approach featured in the main analyses.
